# Characterization of missing values in untargeted MS-based metabolomics data and evaluation of missing data handling strategies

**DOI:** 10.1101/260281

**Authors:** Kieu Trinh Do, Simone Wahl, Johannes Raffler, Sophie Molnos, Michael Laimighofer, Jerzy Adamski, Karsten Suhre, Konstantin Strauch, Annette Peters, Christian Gieger, Claudia Langenberg, Isobel D. Stewart, Fabian J. Theis, Harald Grallert, Gabi Kastenmüller, Jan Krumsiek

**Affiliations:** Institute of Computational Biology, Helmholtz-Zentrum München, Neuherberg, Germany; Institute of Epidemiology II, Helmholtz Zentrum München, German Research Center for Environmental Health, Neuherberg, Germany; Research Unit of Molecular Epidemiology, Helmholtz Zentrum München, German Research Center for Environmental Health, Neuherberg, Germany; German Center for Diabetes Research (DZD e.V.), Neuherberg, Germany; Institute of Bioinformatics and Systems Biology, Helmholtz-Zentrum München, Neuherberg, Germany; Institute of Experimental Genetics, Genome Analysis Center Helmholtz Zentrum München, Neuherberg, Germany; Lehrstuhl für Experimentelle Genetik, Technische Universität München, Freising-Weihenstephan, Germany; German Center for Cardiovascular Disease Research (DZHK e.V.), partner-site Munich, Germany; Department of Physiology and Biophysics, Weill Cornell Medical College in Qatar, Education City, Doha, Qatar; Institute of Genetic Epidemiology, Helmholtz Zentrum München–German Research Center for Environmental Health, Neuherberg, Germany; Chair of Genetic Epidemiology, Institute of Medical Informatics, Biometry and Epidemiology, Ludwig-Maximilians-University, Munich, Germany; MRC Epidemiology Unit, University of Cambridge, Cambridge, United Kingdom; Department of Mathematics, Technische Universität München, Garching, Germany; Institute for Computational Biomedicine, Englander Institute for Precision Medicine, Department of Physiology and Biophysics, Weill Cornell Medicine, New York, USA

**Keywords:** untargeted metabolomics, missing values imputation, limit of detection, batch effects, runday effects, *MICE*, K-nearest neighbor, mass spectrometry

## Abstract

**BACKGROUND:** Untargeted mass spectrometry (MS)-based metabolomics data often contain missing values that reduce statistical power and can introduce bias in epidemiological studies. However, a systematic assessment of the various sources of missing values and strategies to handle these data has received little attention. Missing data can occur systematically, e.g. from run day-dependent effects due to limits of detection (LOD); or it can be random as, for instance, a consequence of sample preparation.

**METHODS:** We investigated patterns of missing data in an MS-based metabolomics experiment of serum samples from the German KORA F4 cohort (n = 1750). We then evaluated 31 imputation methods in a simulation framework and biologically validated the results by applying all imputation approaches to real metabolomics data. We examined the ability of each method to reconstruct biochemical pathways from data-driven correlation networks, and the ability of the method to increase statistical power while preserving the strength of established genetically metabolic quantitative trait loci.

**RESULTS:** Run day-dependent LOD-based missing data accounts for most missing values in the metabolomics dataset. Although multiple imputation by chained equations (*MICE*) performed well in many scenarios, it is computationally and statistically challenging. K-nearest neighbors (*KNN*) imputation on observations with variable pre-selection showed robust performance across all evaluation schemes and is computationally more tractable.

**CONCLUSION:** Missing data in untargeted MS-based metabolomics data occur for various reasons. Based on our results, we recommend that *KNN*-based imputation is performed on observations with variable pre-selection since it showed robust results in all evaluation schemes.

**Key messages:**

- Untargeted MS-based metabolomics data show missing values due to both batch-specific LOD-based and non-LOD-based effects.
- Statistical evaluation of multiple imputation methods was conducted on both simulated and real datasets.
- Biological evaluation on real data assessed the ability of imputation methods to preserve statistical inference of biochemical pathways and correctly estimate effects of genetic variants on metabolite levels.
- *KNN*-based imputation on observations with variable pre-selection and *K* = 10 showed robust performance for all data scenarios across all evaluation schemes.

## Introduction

In epidemiological studies, metabolomics is an established tool that provides insights into disease mechanisms (1), as metabolite profiles generate a molecular readout that is closely linked to the (patho-)phenotype (2,3). Recent metabolomics studies have identified many metabolites as candidate biomarkers for various health conditions, such as diabetes (4–6) and cardiovascular diseases (7,8). Mass spectrometry (MS)-based metabolomics measurements can be performed either in a targeted or untargeted manner (9). In the former, only a limited number of already known and biochemically annotated metabolites are captured. In the latter, the measurements are not limited to predefined signals and offer discovery of novel compounds. While missing values in targeted MS-based data occur rarely, untargeted MS-based techniques typically produce 20-30% missing values, affecting more than 80% of the measured compounds (10–13).

There are various reasons why metabolite concentrations can be missing in an untargeted metabolomics dataset. First, it is possible that the molecules are truly absent from the sample, a situation that may occur e.g. for drug metabolites that only appear in a subset of people taking that medication. On the other hand, there are several technical reasons that could result in missing values, including: (i) instrument sensitivity thresholds, below which concentrations of a specific metabolite might not be detectable in a sample (i.e., below the limit of detection, LOD); (ii) matrix effects that impede the quantification of a metabolite in a sample through other co-eluting compounds and ion suppression; (iii) declining separation ability of the chromatographic column and increasing contamination of the MS instrument; and (iv) limitations in computational processing of spectra, such as poor selection and alignment of the spectral peaks across samples (14).

Commonly, observed patterns of missing data are categorized as either missing completely at random (MCAR), missing at random (MAR), or missing but not at random (MNAR) (15). In the MCAR category, the probability of missing values does not depend on observed or unobserved measurements. In contrast, the occurrence of MAR depends on other observed measurements (for instance, resulting from technical effects, such as overlapping peaks). MNAR describes the occurrence of missing values that depend on unobserved measurements (for instance, due to issues with the performance of the machine).

Although it is clear that the handling of missing values affects all downstream analyses, it is less clear how to appropriately handle their occurrence statistically. A simple *ad hoc* approach is known as complete case analysis (*CCA*), which only considers samples that do not contain any missing values in the metabolites analyzed in each statistical analysis step. However, missing data may occur in some systematic way (i.e., they are dependent on external factors). For example, if all cases in a case-control study have more missing data than the controls, removing observations that are missing will lead to bias in biological interpretation (16). Furthermore, *CCA* can cause severe loss of information and statistical power by excluding a majority of observations if multivariate methods, such as principal component analysis or partial correlation networks, are to be performed.

A widely used and flexible class of missing data strategies is imputation, which involves the replacement of missing values by reasonable substitute values. The most commonly used imputation approaches for metabolomics data assume that missing data occur because they are below the limit of detection (left-censoring, a variant of MNAR). Therefore, all missing entries of a metabolite are replaced by a low constant value, such as the actual LOD (if known), zero, or the smallest value found in the dataset for that metabolite (13). Another LOD-based substitution strategy assumes a parametric left-truncated normal distribution and performs likelihood-based parameter estimation on the observed values to reconstruct the truncated part of the distribution. Missing values are then replaced by numbers drawn from this estimated part (16,17). Additional imputation-based substitution approaches assume MCAR and replace missing values by the mean or median per metabolite (12). Advanced approaches use multivariate statistical methods for imputation, including multiple imputation by chained equations (MICE) (18) and K-nearest neighbors (KNN) imputation (19,20).

Several previous studies have investigated the occurrence and effects of different strategies for missing values in metabolomics data. Taylor *et al.* (21) reported that no single imputation method was universally superior, but constant substitution methods consistently showed poor performance. Gromski *et al.* (12) recommended imputation by Random Forests (RFs) for GC/MS metabolomics data after evaluating the outputs of supervised and unsupervised learning approaches. Di Guida *et al.* (15) investigated various combinations of different preprocessing steps to determine which were the most appropriate for univariate and multivariate analyses of UHPLC-MS metabolomics data. The authors recommended RF and *KNN*-based imputation for PCA and PLS-DA, respectively (15). Armitage *et al.* (10) studied missing values in CE/MS metabolomics data and reported *KNN* imputation to be more effective compared with simpler substitution-based imputation methods. Finally, in a study by Hrydziuszko and Viant (11), a *KNN*-based imputation approach also outperformed competing strategies in an investigation of direct infusion Fourier transform ion cyclotron resonance (DI-FTICR) MS-based metabolomics data.

Despite these advances in our understanding of the effects of imputation on metabolomics data analysis, several aspects have not been addressed by those previous studies. (i) A detailed statistical description of the patterns of missing values in MS-based metabolomics data has not yet been published. Most previous studies evaluated imputation strategies assuming only random or LOD-based missing values without assessing whether this applies to real metabolomics datasets. In particular, the influence of batch effects on the occurrence of missing values has not been investigated in any study. If a cohort comprises a large number of samples, the MS runs usually are spread across multiple days, which is known to influence metabolite measurements due to variation in instrument sensitivity. Here, the LOD itself is also expected to vary across run days, an assumption that has not been explicitly accounted for in any studies. (ii) In addition, a simulation framework that reflects realistic data situations is needed to provide an unbiased evaluation of strategies for handling missing values. Evaluation of previous studies has been biased in the sense that “complete” measured data (created by excluding all variables with missing values) with artificially introduced missing values were simulated, which most likely does not mirror realistic missing value patterns. (iii) Finally, biological validation and biochemical interpretation of the data have not been addressed in the majority of papers. Only Hrydziuszko *et al.* evaluated the ability of different imputation strategies to preserve metabolic differences between biological groups, which then were related to KEGG pathways (11).

In the present study, we analyzed patterns of missing data and evaluated the performance of various imputation strategies for untargeted MS-based metabolomics data from serum samples of the German Cooperative Health Research in the Region of Augsburg (KORA) F4 cohort. Data were measured on a typical, widely used untargeted MS-based metabolomics platform (Metabolon, Inc., USA) and should be representative of many untargeted population-scale metabolomics studies. The study consisted of three steps: (i) We described and analyzed patterns of missing values and their possible underlying mechanisms in a real untargeted metabolomics dataset. In particular, we investigated the occurrence of missing values within and across batches of measurements. (ii) The insights gained from these analyses were used to introduce realistic patterns of missing data into simulated data. We applied 31 imputation methods to the datasets and evaluated them with respect to their ability to achieve correct statistical estimates and hypothesis test results in various data scenarios. (iii) Finally, the imputation methods were applied to real metabolomics data (KORA F4), followed by two biologically-driven evaluation schemes. First, we assessed how accurately real biochemical pathways were reconstructed in data-driven correlation networks inferred from the imputed data. Second, we verified whether imputation led to a gain in statistical power, while preserving effects of genetic variants on metabolite levels. The study workflow is visualized in Figure 1.

**Figure 1.**
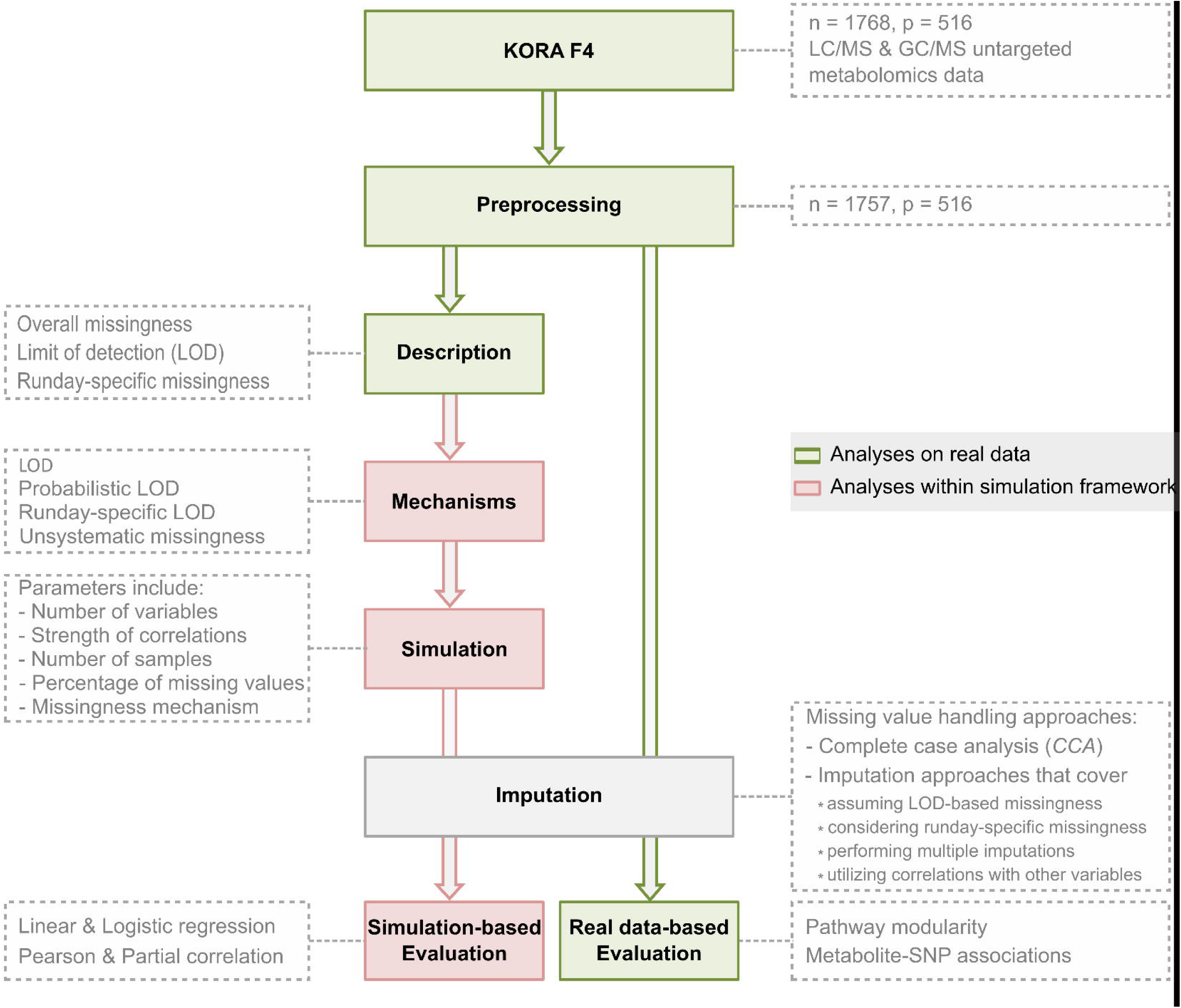
Flow chart of the study design. Pre-processed KORA F4 metabolomics data were used to analyze patterns of missing values in the dataset. Possible underlying mechanisms were inferred and implemented in a simulation framework to generate data resembling the observed patterns. Based on these simulated data, imputation methods with different characteristics were applied and evaluated. Finally, the same imputation approaches were evaluated using KORA F4 metabolomics and genomics data.

## Results

### Characterization of missing data patterns in KORA F4 untargeted metabolomics data

We used an untargeted metabolomics dataset from the KORA F4 study, which was generated from fasting serum samples measured on three platforms: LC/MS in both positive (LC/MS+) and negative modes (LC/MS–), as well as a GC/MS platform. After log-transformation and outlier handling (see Methods), 1757 samples and 516 metabolites were available for analysis.

The dataset contained 19.41% missing values, with 416 (80.6%) metabolites and all observations showing at least one missing value. The majority (301) of these 416 metabolites had fewer than 10% missing values (Figure 2A). For only 9.9% (51) of the metabolites, more than 70% of the measurements were missing. The amount of missing values per observation ranged from 11.4% to 32.2%, with an average of 19.6% (Figure 2B).

**Figure 2.**
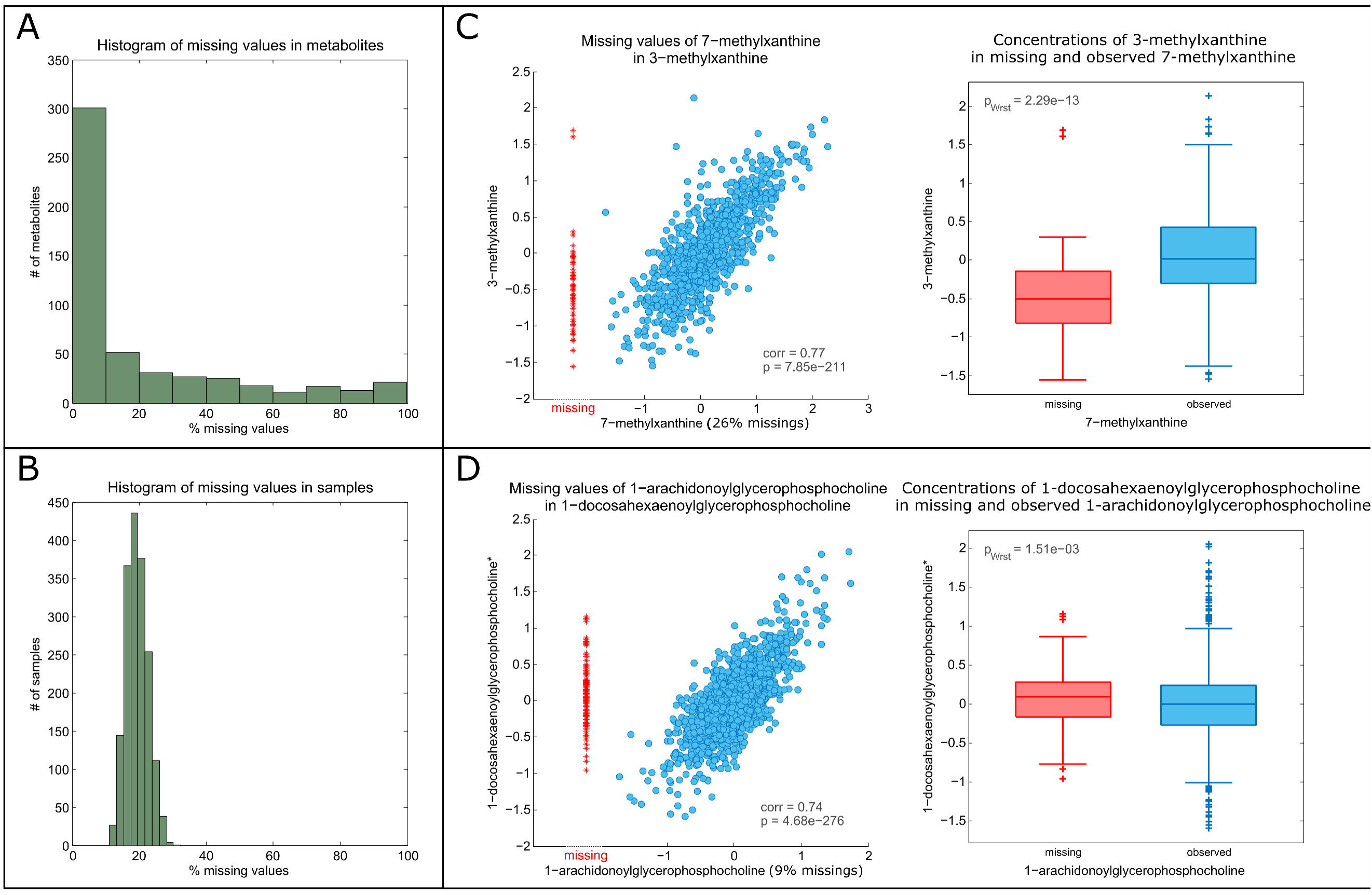
Overall amounts of missing data and LOD effects. (A,B) The overall fraction of missing values across metabolites and observations, respectively. (C,D) Scatter plots and boxplots of selected metabolite pairs to illustrate missing data due to LOD and non-LOD effects, respectively. Blue observed concentrations. Red - observed values of the auxiliary metabolite in observations with missing values of the investigated metabolite. Note that red data points are not part of the x-axis but were plotted in the same scatterplot for clarity. *corr*= correlation, *p* = p-value of correlation, ***p***_***Wst***_ = p-value of Wilcoxon–Mann–Whitney test.

#### LOD-based missing values

For metabolomics data, a common assumption is that missing values occur because of low concentrations that are below the limit of detection. To explore this assumption, we analyzed missing values of a metabolite using a second, strongly correlated metabolite, which we term the *auxiliary* metabolite. The auxiliary metabolite is defined as the metabolite with the highest correlation (*r*) to the given metabolite. Due to its strong correlation, we assume that insights into the pattern of missing values of a metabolite can be gained from the corresponding non-missing observations of its auxiliary metabolite. For example, assuming that metabolite A has missing values in certain observations for which its auxiliary metabolite B has measurements. If these measurements in B are low then a missing value in A most likely occurred because the actual concentrations were below the LOD. We required a minimum correlation of *r* = 0.3 for auxiliary metabolites, but other values gave qualitatively similar results (File S1).

Overall, an auxiliary metabolite was available for 56.6% of the metabolites. Of those, 62.0% showed a clear tendency for missing values to below the LOD (see Methods and File S1). An example for a clear LOD-tendency is shown for 7-methylxanthine in Figure 2C. This compound is a metabolite of caffeine metabolism that is correlated with 3-methylxanthine. The majority of observations with missing data in 7-methylxanthine showed low values for 3-methylxanthine, indicating that the 7-methylxanthine values were most probably below the LOD. An example for a metabolite pair that does not show an LOD-based missingness pattern is provided in Figure 2D for 1-arachidonoylglycerophosphocholine (1-AGPC) and its auxiliary metabolite 1-docosahexaenoylglycerophosphocholine (1-DGPC). Unlike the previous example, observations with missing data for 1-AGPC showed values varying over the whole range of 1-DGPC. Consequently, this suggests that LOD does not adequately explain the pattern of missing values for 1-AGPC. Scatterplots of investigated metabolites and their corresponding auxiliary metabolites, as well as boxplots of concentrations in the auxiliary metabolites for missing and non-missing observations in the investigated metabolites can be found in File S1.

Although the LOD-tendency was observed for many metabolites, there was no clear LOD threshold separating missing and observed measurements across all metabolites (Figure 2C), which would have been the case if LOD was the only underlying mechanism for missing data. Instead, the values of the auxiliary metabolites with missing values in the investigated metabolites were spread broadly over a range of lower values, indicating a blurred rather than a single fixed LOD for all metabolites.

#### Run day-dependent missing values

Batch (run day) effects also can drive systematic patterns of missing data due to daily variation in instrument sensitivity. To examine whether missing data depended on overall run day quality, we examined the amount of missing values per run day for each platform (LC/MS+, LC/MS–, or GC/MS). Subsequently, we investigated whether metabolites were affected differently by runday quality.

The KORA F4 samples were measured on 53 run days with 34 samples on average per day. If missing values were dependent on run day quality due to variation in instrument performance (e.g., caused by LC or GC column decline), we would expect there to be some days for which samples overall contained more (“bad” run day) or fewer (“good” run day) missing values compared with the average. Indeed, we observed such “bad” and “good” run days for all three platforms (Figure 3A). While the run day-specific amount of missing values tended to be correlated between LC/MS– and LC/MS+ (correlation of the run day-specific median of missing values between the two platforms was *r* = 0.36), there was no correlation between LC/MS+/– and GC/MS. This suggests that changes in instrument performance, rather than global effects (such as those that could originate from sample preparation) were responsible for differences in run day quality.

**Figure 3.**
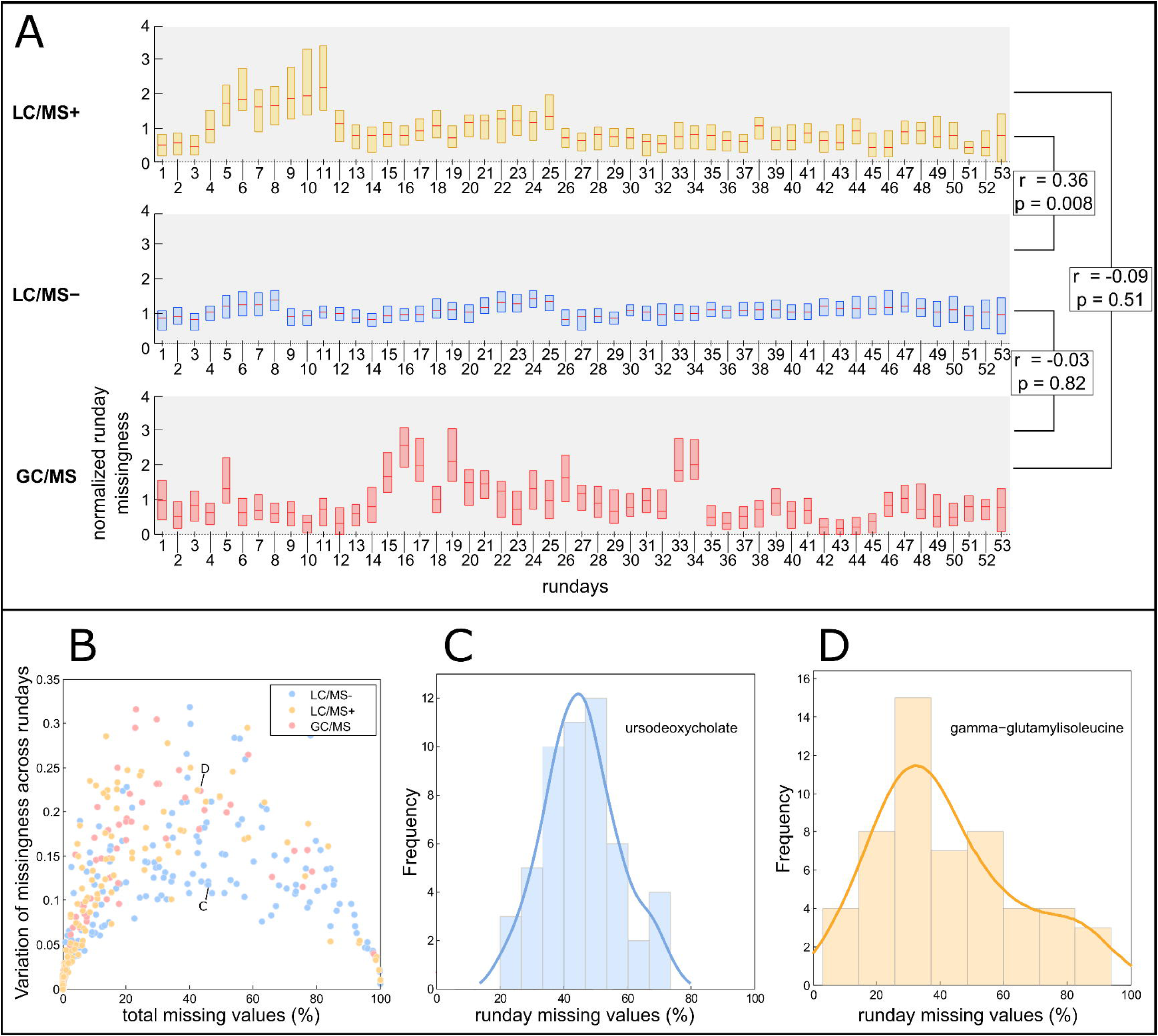
Run day-dependent effects on missing data. (A) Normalized amount of missing values per run day in each platform (LC/MS+, LC/MS-, GC/MS). For a given metabolite and run day, the normalized amount of missing data per run day was calculated as the number of missing values for the respective metabolite on the respective run day divided by the total number of observations for that run day, divided by the median amount of missing data of that metabolite over all run days. Thus, a normalized run day-missingness of 1 is the average run day-missingness for a given metabolite. Pearson correlation coefficients were calculated across all pairs of platforms. (B) Standard deviation of missing values across run days, depending on the total amount of missing data for each platform. Each dot in the plot shows the total proportion of missing values and the run day variation for one metabolite. (C)–(D) The distribution of the total amount of missing values is shown for a metabolite with moderate (ursodeoxycholate) and high (gamma-glutamylisoleucine) standard deviation.

Although there was an overall effect of run day quality on the pattern of missing values, we observed considerable differences in the standard deviations (SD) of run day-specific missing values for metabolites with the same amount of missing data (Figure 3B). This suggests that metabolites were affected differently by run day quality. For example, the bile acid ursodeoxycholate (46% total missing data) showed relatively low variation in run day missing data (SD = 0.12) (Figure 3**Figure 3**C). However, for gamma-glutamylisoleucine (Figure 3D), a metabolite with a similar total amount of missing values (42%), the observed variation in missing data across run days was substantially larger (SD = 0.22).

#### Run day-dependent LOD mechanism

The observed run day-dependent pattern of missing data, together with the blurred LOD-based pattern, suggests that different run days may exhibit different LODs, which contributed to the blurred global LOD effect. To verify this, we calculated the correlation between run day mean and run day missingness for all metabolites. A histogram of the correlation coefficients is shown in Figure 4A. The majority of metabolites displayed a strong tendency for negative correlations. An example for run day-specific LODs is shown in Figure 4B–C: for 7-methylxanthine, the correlation of run day mean and the run day-specific amount of missing values is *r* = −0.68 (Figure 4B). Run days with low means tended to have a higher amount of missing values (Figure 4C). Density plots for all metabolites before and after run day normalization can be found in File S2.

**Figure 4.**
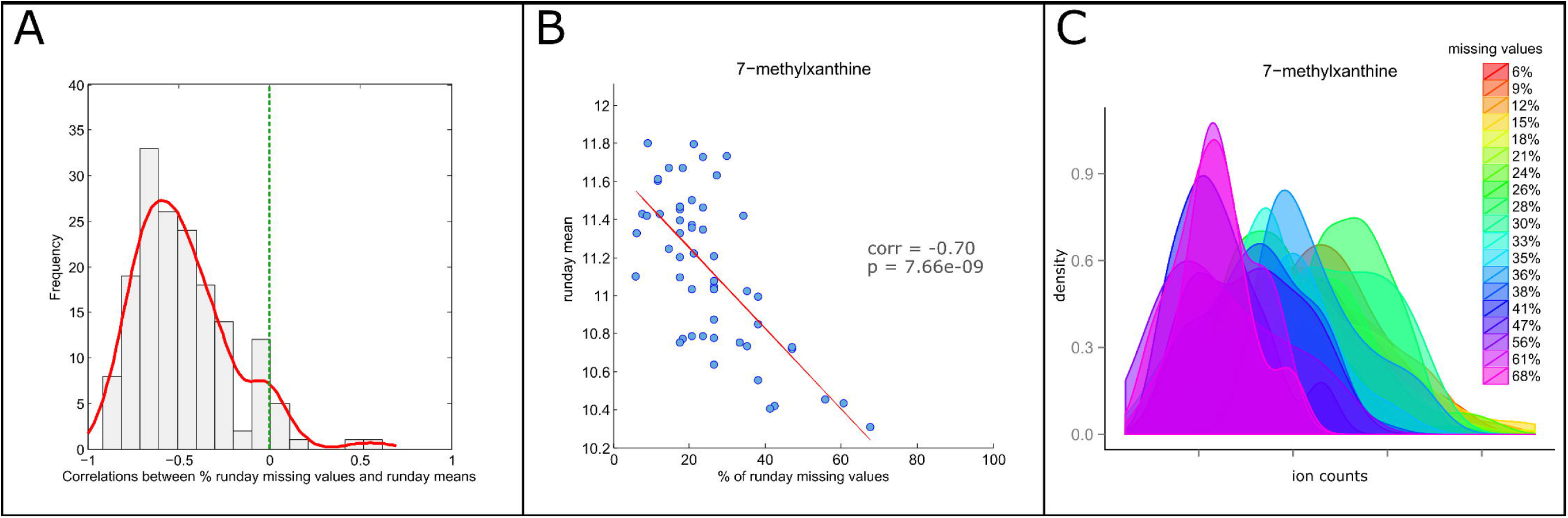
Run day-dependent LOD. (A) Histogram of Pearson correlation coefficients of the percent of missing values and run day means. (B) Scatterplot of run day mean versus percent missing values, with 7-methylxanthine as an example of a negative correlation. (C) Run day distributions of 7-methylxanthine before run day normalization.

Taken together, we observed that batch (run day) effects on the limit of detection can result in a blurred LOD-effect after run day normalization, which can explain patterns of missing values in most, but not all, metabolites.

### Evaluation of imputation approaches in a simulation framework

As shown in the previous analyses, not all of the missing data in MS-based metabolomics studies can be attributed to run day-dependent LOD-based missing data. Thus, the optimal imputation approach should perform well across all possible patterns. We conducted a simulation study to compare statistical estimates between imputed and complete data. We simulated incomplete data according to the patterns of missing values observed in the real metabolomics data and imputed these data using various imputation approaches. We then evaluated these approaches for recovering correct statistical estimates after conducting correlation and regression analyses.

#### Simulation setup and evaluation criteria

We simulated six mechanisms for missing data derived from observations in the real data (see Methods, File S3, and Figure 5A–E): (i) *Fixed LOD,* as an extreme form of systematic missing values below a global LOD; (ii) *Probabilistic LOD*, where the probability of a missing value increases at lower values, which should resemble the blurred LOD-based patterns observed in the real data; (iii) *Run day-specific fixed LOD*, where LOD is assumed to vary across run days; (iv) *Run day-specific probabilistic LOD*, where a probabilistic form of LOD is assumed to occur across run days; (v) *Unsystematic (random) missingness*, for missing data with an unknown reason; and (vi) *Mixtures of LOD-based and unsystematic missingness*. Based on these 6 mechanisms, we created various parameter scenarios resembling realistic conditions. For each scenario, we conducted 250 simulations to assess whether the imputation methods could reconstruct statistical estimates of Pearson correlation, partial correlation, linear regression (results shown in File S3), and logistic regression. To this end, we calculated type 1 error as the proportion of simulations in which a significant estimate was obtained when the true correlation was equal to zero. In addition, we calculated power as the proportion of significant estimates when the true correlation was unequal to zero. We also estimated bias, which is shown in File S3. A detailed description of the simulation and evaluation framework is also provided in File S3.

**Figure 5.**
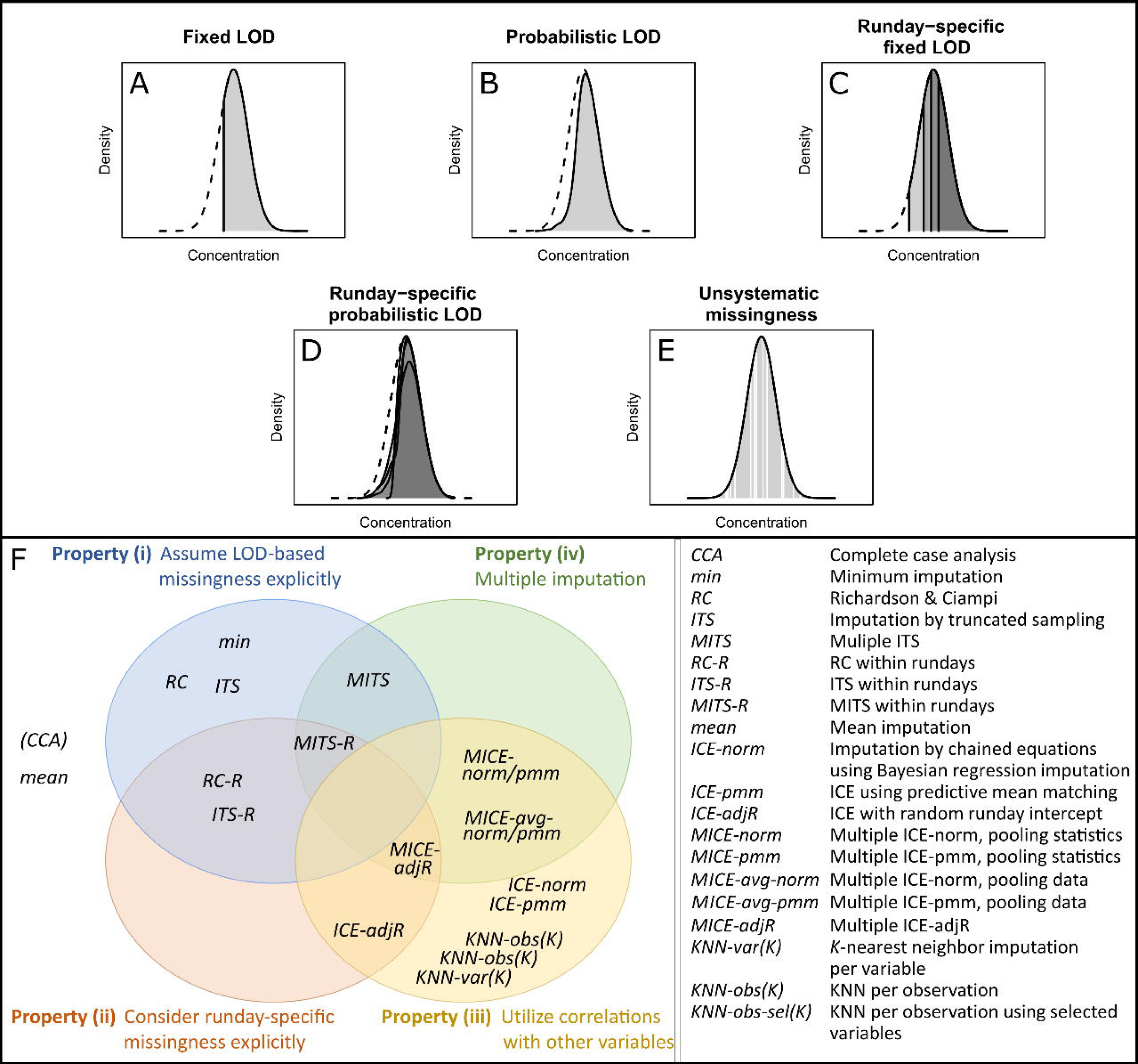
Mechanisms of missing data and imputation approaches used in the simulation study. (A)–(E) Mechanisms of missing values used in the simulation study, based on evidence from real metabolomics data. (F) Venn diagram of imputation methods showing different characteristics. Note that the figure contains complete case analysis (*CCA*), which is not an imputation method, and is noted in brackets. *CCA* and *mean* were placed outside the Venn diagram, as they do not comprise any of the four characteristics. LOD: limit of detection.

#### Missing data handling strategies

We applied 31 imputation approaches (see Figure 5F; detailed descriptions in Methods and File S4) on the simulated data. Some were adapted to account for run day-specific missing values. The imputation approaches followed different concepts, which could have one of the following four properties or combinations thereof: (i) approaches that explicitly assume LOD-based missing values, (ii) approaches that consider run day-specific missing values, (iii) multivariate procedures using correlations among variables, and (iv) multiple imputation (MI) strategies. The MI approaches usually comprise imputation, analysis, and pooling steps. In the first step, the incomplete data are imputed *m* times to produce *m* complete datasets. Subsequently, statistical analysis is performed on each of the *m* complete datasets and then the *m* analyses are combined to one final result.

#### Simulation results

In the following, we evaluate the performance of the four imputation properties (i)–(iv) introduced above. Simulation results from other data scenarios, all variations of the imputation approaches used, and the combination of parameter settings are available in File S5.

Property (i): Methods that explicitly assume LOD-based missing values and perform imputation globally without taking run day information into account (*min*, Richardson & Ciampi (*RC*), imputation by truncated sampling (*ITS*)), showed inflated type 1 error rates and low power for both correlation and regression analysis. This was expected for three reasons. First, for a data scenario with run day-dependent probabilistic LOD-based missing values, these methods underestimate the LOD for most of the rundays and replace missing entries by too low values (Figure 6A). Second, for a data scenario with random missing values, they expectedly fail since the underlying assumption of an LOD is not met (Figure 6B). Finally, *min* and *RC* impute a metabolite by replacing all of its missing entries by a constant value, which substantially distorts the metabolite distribution (see File S5).

**Figure 6.**
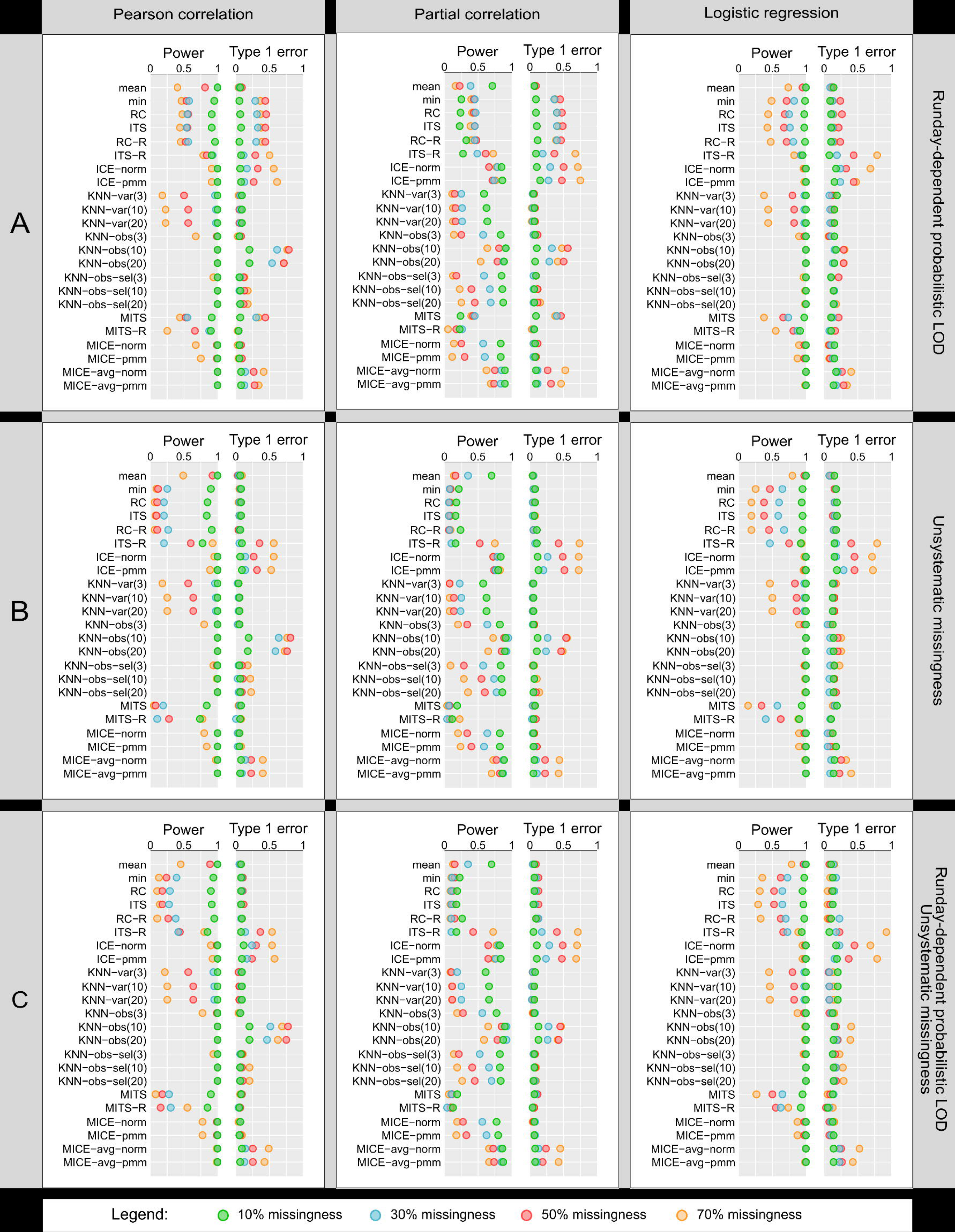
Simulation results for Pearson, partial correlation, and logistic regression analysis. Performance of imputation approaches in data scenarios where (A) both variables followed a run day-specific probabilistic LOD mechanism, (B) both variables showed non-systematic patterns of missing data, and (C) one variable with run day-specific probabilistic LOD-based missing data and the other variable showed non-systematic patterns of missing data. Type 1 error and power reflect the false positive and true positive rate of hypothesis testing, respectively. Note that power = 1 - type 2 error rate. Note further that due to readability issues, only KNN-based imputation methods with *K* = 3, 10, and 20 were included, whereas KNN imputation with *K* = 1 and 5 can be found in File S5.

Property (ii): The LOD-based methods that take run days into account (*RC-R, ITS-R*) were expected to perform well in a simulated data scenario with run day effects (Figure 6A). Unexpectedly, we observed an inflated type 1 error rate and decreased power for all three statistical analyses (Pearson correlation, partial correlation, and logistic regression). *RC-R* and *ITS-R* assume that the observed values of a metabolite follow a truncated normal distribution, which is parametrized by maximum likelihood estimation (MLE), in order to replace missing values with randomly drawn values from the truncated part. The instability of MLE due to small sample sizes available within run days could explain the poor performance of these approaches. The same poor performance was observed for scenarios with a mixture of run day-dependent LOD-based and random missing values (Figure 6C). For the dataset with only random missing values, LOD- or run day-based approaches showed the expected strong reduction in power since here the underlying assumption of a truncated normal distribution is false (Figure 6B).

Property (iii): Multivariate approaches (imputation based on chained equations *(ICE)* and *KNN*-based imputation) take into consideration the correlation between variables or observations. *ICE* approaches had high power, but an increased type 1 error rate when missing value proportions increased (Figure 6). *KNN*-based imputation on observations with variable pre-selection and *K* = 10 (*KNN-obs-sel(10)*) was one of the best performing methods with high power and an overall marginal type 1 error rate, even for a high amount of missing values. The power for *KNN-obs* was also high, but it showed high type 1 error rate and therefore a poor ability to correctly identify truly absent associations. In contrast, *KNN-vars* had a low type 1 error rate, but decreased power, which became more pronounced at higher amounts of missing values.

Property (iv): Single imputation procedures often underestimate the variability of statistical estimates, resulting in inflated type 1 error rates. This should be avoided by approaches performing multiple imputations (MI). MI versions based on LOD-(*MITS*) and run day-effects (*MITS-R*) indeed had decreased type 1 error rates, although power was low (Figure 6). *MICE* with Bayesian linear regression (*MICE-norm*) or predictive mean matching (*MICE-pmm*) as imputation model showed negligible type 1 error rates and high power for all scenarios with up to 50% missing values. At higher amounts of missing data, the power decreased considerably, but the type 1 error remained marginal (File S5). A slight modification of the *MICE* algorithm applied widely in the metabolomics field (here termed *MICE-avg*) was performed on each imputed data, and comprised the pooling of the imputed data with subsequent statistical analyses rather than pooling the statistical estimates after analysis. This approach showed high power, but increased type 1 error rates, in particular for >30% missing values.

Taken together, when considering all patterns of missing data and all evaluation criteria, *KNN-obs-sel(10)* and *MICE-norm* were the most robust approaches. For higher amounts of missing data (≥50%), *MICE* showed a strong decrease in power with marginal type 1 error, whereas *KNN-obs-sel(10)* had only slightly increased type 1 error rates with high power.

### Evaluation of imputation approaches on real MS-based metabolomics data

We conducted a biological evaluation of all approaches using the metabolomics data from the KORA F4 population study. An objective criterion for evaluation is challenging to construct, since the true values underlying the missing ones are unknown. We devised two indirect tests that assessed imputed values for biological validity. First, we assessed the ability of imputation methods to statistically reconstruct biochemical pathways in metabolomics data. Second, we evaluated the gain in statistical power while preserving the true effect size of genetic variants (SNPs) on metabolite levels.

#### Evaluation based on pathway modularity

GGMs are based on partial correlations and reflect conditional dependencies in multivariate Gaussian distributions (5,22). When applied to metabolomics data, they reconstruct a precise picture of the metabolic network, showing a modular topology with respect to known pathways. In other words, metabolites will tend to be correlated with other metabolites from the same biochemical pathway (5,22,23). We used this pathway-based modularity in a metabolic network as a quality criterion to indicate whether the imputation methods generally were capable of maintaining biochemically valid edges.

Each imputation strategy was applied to the KORA F4 metabolomics data, and a GGM was estimated for each obtained dataset. Subsequently, we used *a priori* pathway annotations from Metabolon Inc., where each metabolite was assigned to one pathway (e.g., branched-chain amino acids, lysolipids, xanthines) to calculate pathway-based modularity (*Q*), according to (22,24). This measure reflects the ratio of metabolite correlations within *versus* across pathways. A high *Q* value indicates a dense within-pathway correlation compared with cross-pathways. Variability was estimated by bootstrap resampling (see Methods).

Across all datasets, we obtained modularity values ranging from 0.384 to 0.434 (Figure 7A). Imputation methods that explicitly considered the LOD-based mechanism and their run day-specific versions (Figure 5, property (ii)) did not outperform alternative approaches. Multivariate, single imputation methods (property (iii)) yielded low *Q* values, except for *KNN-obs-sel*, which achieved the overall third best result (*Q* = 0.422 for *K* = 10) (Figure 5). The performance of *KNN*-based imputation methods strongly depended on the definition of neighbors (variables or observations) and on the number of these neighbors (*K*). The MI procedures (property (iv)) *MITS*, *MITS-R*, and *MICE-avg* performed poorly, whereas the networks generated on *MICE* imputed data showed the overall highest modularity (*Q* = 0.434 and *Q* = 0.424 for *MICE-norm* and *MICE-pmm*, respectively) (Figure 5). Overall, the three best performing approaches were *MICE-norm*, *MICE-pmm*, and *KNN-obs-sel(10)*.

**Figure 7.**
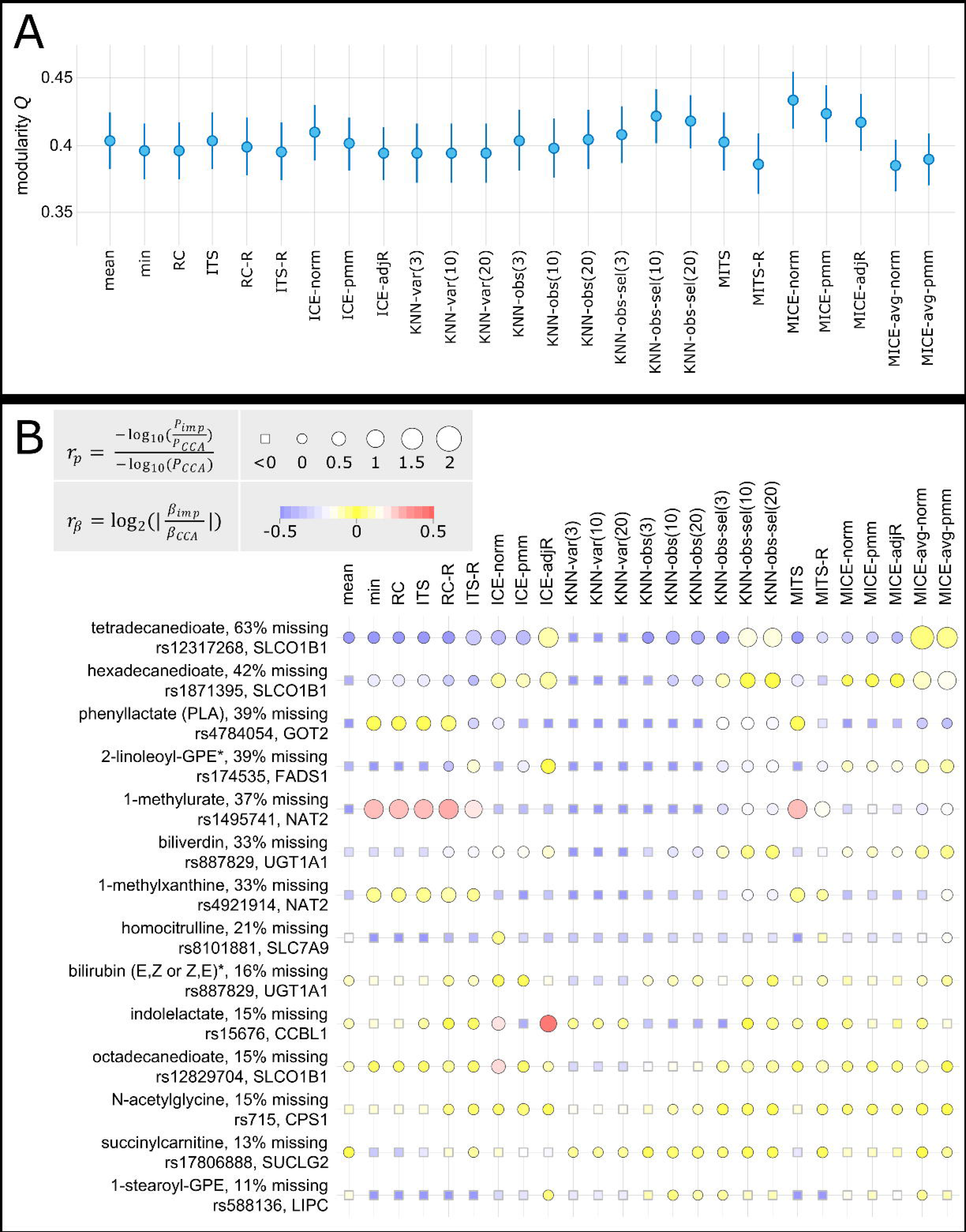
Evaluation of imputation approaches on real data. (A) Pathway-based modularity for each imputation strategy. Modularity *Q* was calculated based on pathways. Vertical lines represent bootstrap-based confidence intervals (1000 times resampling). (B) The ability to gain statistical power and to preserve real metabolite-SNP associations after imputation. Circle color represents the ability of imputation methods to preserve effect sizes, with red and blue indicating possible overestimation and underestimation, respectively, and yellow corresponding to cases with good preservation of the association. Circle size depicts the gain in statistical power after imputation. The bigger the circle the higher the statistical power gain after imputation compared to *CCA*. Squares correspond to cases where no statistical power was gained. Note that due to readability issues, only KNN-based imputation methods with *K* = 3, 10, and 20 were included, whereas KNN imputation with *K* = 1 and 5 can be found in File S6 and Table S8.

#### Evaluation based on metabolite-SNP associations

Using KORA F4 data (n = 1750), we determined the ability of imputation methods to gain statistical power compared with complete case analysis (*CCA*, deleting samples with any missing values) while preserving the effect of genetic variants on metabolite levels in human blood. For the evaluation, we selected a set of metabolite-SNP associations from a previous genome wide association study (GWAS) in the KORA F4 and TwinsUK cohorts, for which a functional connection between the gene and the metabolite was biologically evident (Table S8) (25). For example, GOT2 (*rs4784054*), which was associated with concentrations of phenyllactate, encoded an enzyme that catalyzes the conversion of phenylalanine to phenylpyruvate, which is then converted to phenyllactate (25,26).

We investigated the gain in statistical power when using imputed datasets compared with the power obtained with *CCA* for 18 of such metabolite-SNP pairs, where the metabolite had between 10% and 70% missing values. Statistical power gain was calculated as the negative log10 of the ratio of the p-values estimated for the imputed data to the p-values estimated for *CCA* in corresponding linear regression models (detailed results in File S8 and Table S8). A high ratio indicates greater power for imputed data. As a second evaluation criterion, we calculated the log2 absolute ratio of the effect sizes obtained from the regression models for imputed data and those derived from *CCA* in KORA F4 (see Methods). A log2 ratio close to zero indicates that the imputation method was able to preserve effect sizes, whereas imputations yielding a highly negative or positive log2 ratios indicate underestimation or overestimation of the effect sizes, respectively.

Imputation with LOD-based methods (property (i)) yielded a gain in power for up to seven genetic associations of the 14 metabolites (Figure 7**Figure 7**). For two of these associations (tetradecanedioate and SLCO1B1; and hexadecanedioate and SLCO1B1), effect sizes were underestimated, and for the association between 1-methylurate and NAT2, the effect size was overestimated across all methods, except for *MITS-R*. Run day-specific imputation methods (property (ii)) performed well, with *ITS-R* yielding the highest number of associations (12) with greater statistical power, of which seven showed effect sizes similar to effect sizes derived from *CCA*. The best methods among multivariate approaches (property (iii) and (iv)) were *MICE-avg-norm*, *KNN-obs-sel(10)*, *and KNN-obs-sel(20)*, all three of which generated a gain in statistical power for 12 associations. These methods also showed good performance in preserving genetic effects and did not show severe overestimation or underestimation of effect sizes. *MICE-norm/-pmm/-adjR* showed only moderate performance with a power gain for seven associations.

In an additional analysis, we used results from the EPIC-Norfolk cohort with n = 10 634 subjects (27), to assess the ability of imputation methods to preserve effects of genetic variants on metabolites. We hypothesized that the effect sizes would be estimated more accurately in this much larger dataset, and effect sizes obtained with KORA F4 imputed data should approximate effect sizes derived from EPIC-Norfolk. Overall, we observed that the majority of SNP-metabolite pairs showed either an overestimation or an underestimation of effect sizes across all imputation methods. This tendency might reflect differences between the cohorts KORA F4 and EPIC-Norfolk rather than differences between imputation strategies (see detailed results in File S7 and Table S8).

Overall, for nearly all metabolite-SNP pairs, this analysis showed that statistical power was increased by imputing missing values and the effect sizes could be preserved. *ITS-R*, *MICE-avg-pmm, KNN-obs-sel* with *K* = 10 and *K* = 20 were the imputation methods that generated the highest number of associations (12) and resulted in a gain in statistical power compared with *CCA*.

## Discussion

In this study, we investigated patterns of missing data in a typical example of untargeted MS-based metabolomics data and their possible underlying mechanisms. Insights gained from these analyses were used to generate simulated data that reflected the real data situation for a comprehensive evaluation of 31 imputation methods. Finally, we applied the imputation strategies to real MS-based metabolomics data from the German KORA F4 study and evaluated them using biological validity measures.

For metabolomics data, an intuitive assumption is that missing data occur when metabolite concentrations fall below the machine’s LOD. Indeed, we found evidence for systematic patterns of missing data due to LOD- and batch-effects for a large proportion of the analyzed metabolites. Missing data were found to be influenced by run day quality, although metabolites varied in their susceptibility to this effect. Finally, we found a negative correlation between run day mean and missing data per run day, further confirming LOD-based mechanism within run days. The existence of multiple run day-dependent LODs possibly accounted for the blurred rather than fixed global LOD observed in the data. It has been suspected that multiple detection limits arise from factors such as batch (run day) effects (27). However, to the best of our knowledge, this is the first time that these effects have been systematically explored so far.

We evaluated 31 imputation methods in an evaluation framework consisting of three schemes: (i) unbiased estimation of statistical estimates and hypothesis test results based on simulated data, (ii) statistical reconstruction of biochemical pathways in metabolic networks, and (iii) the ability to preserve effects of genetic variants on metabolite levels while allowing for a gain in statistical power.

*MICE-norm* was the best performing imputation method for evaluation scheme (i) and (ii), but it showed only moderate performances in the metabolite-SNP analysis. One major drawback of this method is that multiple imputations have to be performed, making these approaches statistically and computationally challenging. For *m* imputations, the desired statistical analyses must be performed on each of the *m* imputed datasets, and then the resulting *m* estimates must be combined to one statistical result. A widely applied alternative is to perform *m* multiple imputations and then combine the *m* complete datasets to one final dataset containing the average of the imputed values (*MICE-avg*). That is, *MICE-avg* does not require statistical estimates to be pooled, and therefore, it is much easier to apply. However, this simplicity is accompanied by an underestimation of metabolites’ variances, resulting in poorer performance of statistical estimation (correlation and regression coefficients) and reconstruction of biochemical pathways.

A feasible, but better performing method was *KNN-obs-sel(10)*, which uses *KNN-*based imputation on observations with variable pre-selection and *K* = 10. This method ranked highly in all evaluation schemes. Other *KNN*-based imputation schemes, including *KNN*-based imputation on variables (*KNN-vars*) and on observations without variable pre-selection (*KNN-obs*), consistently showed poor performance across all evaluation schemes. Our results are in line with observations from previous studies, where *KNN*-based imputation performed well (10,11,15,28). However, we also observed that variations of *KNN* imputation lead to substantially different results, as in previous studies (20,28).

Although we observed LOD- and run day-based effects in real metabolomics data, methods that explicitly consider this information did not outperform competing approaches in the first two evaluation schemes. This is likely due to the fact that they perform imputation in a univariate manner without taking the correlation between the variables into account. Moreover, all of these LOD-based methods include maximum likelihood estimation in their imputation process, which was found to perform well only for larger sample sizes in previous studies (27,29). In our study, the number of observations within run days is limited, resulting in considerable instability of the MLE. LOD-based run day-dependent methods performed well with respect to gain in statistical power in the analysis of metabolites–SNP associations.

In summary, we have presented a detailed description of patterns of missing data in untargeted MS-based metabolomics data. In particular, we considered, for the first time, the effects of run days on systematic patterns of missing data. Our work showed that missing data occur in most cases due to LOD effects, which are moreover run day-dependent. Nevertheless, *MICE* and *KNN*-based imputation, methods that do not explicitly consider LOD-based effects, performed best when tested in both statistical and biological evaluation schemes. This is most likely because these methods take into account multivariate dependencies within the data. The two approaches are For future studies, we recommend *KNN-*based imputation on observations with *K* = 10, since it consistently performed well across all data scenarios and all evaluation schemes, and is computationally non-demanding for daily data analysis.

## Material and Methods

### Study cohort, metabolomics and genotype measurements

Data from 1768 fasting serum samples of the German Cooperative Health Research in the Region of Augsburg (KORA F4) population cohort (30) was used, comprising 910 females and 858 males. Age distribution was 60.53 ± 8.79 years for females and 61.20 ± 8.78 years for males. Body mass index (BMI) distribution was 27.88 ± 5.24 kg/m^2^ for females and 28.46 ± 4.29 kg/m^2^ for males.

Serum metabolomics measurements were performed on three platforms, LC/MS– (negative mode), LC/MS+ (positive mode), and GC/MS by Metabolon, Inc. (Durham, NC, USA). The 1768 serum samples were measured on 53 different run days, with 34 samples on average per run day. A total of 516 metabolites were quantified, of which 303 had an identified chemical structure. A more detailed description of sample acquisition, experimental procedures, and metabolite identification can be found in File S10.

Each known metabolite was annotated with one of 68 pathways by Metabolon, Inc. A full list of all measured metabolites, including pathway annotations, can be found in Table S9. For correlation analysis, data were normalized for run day-effects by dividing each metabolite by run day median. Since metabolite measurements were assumed to follow a log-normal distribution, the data were log-transformed for all statistical analyses. The run day-corrected and log-transformed data were used to determine outlier samples. Eleven individuals with a Mahalanobis distance (calculated across the complete dataset) greater than four SD from the mean were considered outliers and excluded from the dataset. For the biological evaluation schemes, age, sex, and BMI were used as standard covariates. Seven samples were excluded due to incomplete information in these phenotypes, resulting in 1750 individuals in total.

The KORA F4 cohort was genotyped using the Affymetrix Axiom platform. After quality control, genotype data (measured or imputed according to data from the 1000 genomes project, phase 1 version 3) were available for 1685 of the 1750 individuals.

### Missing data in KORA F4

To explore the mechanism for the missing data of a given metabolite *m*, a second (auxiliary) metabolite *m*_*aux*_ was used. *m*_*aux*_ was defined as the metabolite with the strongest Pearson correlation to *m* (at least 0.3). An LOD-tendency was assumed if the average value of *m*_*aux*_ in samples with missing values in *m* was significantly lower than the average of *m*_*aux*_ in samples with measured values in *m*. Significance was assessed using Wilcoxon–Mann–Whitney tests with *α* = 0.05 after Bonferroni correction for multiple testing.

For all correlation analyses, only metabolites with more than 10% and less than 70% overall missing values were considered.

In order to explore whether missing values varied among run days, the normalized proportions of missing values among the 53 run days were compared within each platform. For a metabolite *m* and a run day *d*, the normalized amount of run day-specific missing values was calculated as the number of missing values for *m* in *d* divided by the total number of samples measured in *d*, divided by the median value of missing data of *m* over all run days.

### Simulation study

Insights gained from the analyses of missing values in real MS-based metabolomics data were used to create artificial data that best mirror reflected patterns of missing data. A brief overview of the simulation framework is provided below, and a detailed description can be found in File S3. For each set of parameters corresponding to a certain data situation, 250 random datasets were generated. For each dataset, two variables were simulated by drawing from a multivariate normal distribution, with sample sizes ranging from 100 to 1000, and with means equal to zero and covariance chosen such that variances were equal to one (representing scaled variables). The Pearson correlation between the two variables was ranged from 0 to 0.4. In addition, for the multivariate analyses and to evaluate imputation methods that apply to a multivariate strategy, auxiliary variables correlated with the two main variables were introduced. Their number and correlation strength were chosen to match the real data (for details, see File S3).

Simulated observations were randomly assigned to “run days” with the number of run days chosen such that each run day comprised 34 observations, according to the average number found for the real KORA F4 measurements.

A proportion of missing values (10%, 30%, 50%, and 70%) was introduced into the main variable pair according to different mechanisms derived from our observations in the KORA F4 Metabolon data (Figure 5, File S3).

We used the following parameter settings for the results in the main manuscript: moderate variability of missing data across run days (see File S3), uncorrelated run day-specific missing patterns of the metabolite pair, and varying association of the inverse relation between metabolite concentration and missing values, at *n* = 250 and in the presence of informative auxiliary metabolites. For Pearson and partial correlation analysis, both main variables had the same degree of missing data. For logistic regression analysis, the predictor variable had a mixture of 50% run day-dependent probabilistic LOD-based missing data and 50% non-systematic missing data. Results for more parameter settings can be found in File S5.

### Imputation approaches

A variety of imputation methods (Figure 5**Figure 5**) were selected because they were reported in the context of metabolomics data or were developed and adopted to address characteristics in the current dataset.

***Mean imputation (mean):*** All missing values of each incomplete variable are replaced by the average of the observed values of that metabolite. ***Minimum imputation (min):*** All missing values of each incomplete variable are replaced by the smallest observed value of that metabolite (5,13,16). ***Richardson* & *Ciampi (RC):*** Assuming that missing values occur due to LOD and the observed metabolite values follow a left-truncated normal distribution, maximum likelihood is used to estimate this distribution. A missing value *x* is then replaced by the expected value of *x* conditional on *x* being below the LOD, *E*(*x*|*x* ≤ *LOD*) (17). ***Imputation by truncated sampling (ITS):*** This is an extension of the *RC* method, where the missing values are replaced by randomly drawn values from the censored part of the estimated truncated normal distribution. ***Multiple imputation by truncated sampling (MITS):*** *ITS* is applied as described above, but multiple imputation is performed according to Rubin’s rules (31) using the *R* package *mice*, version 2.25. These rules include: (i) the datasets are imputed m times, (ii) each of the *m* completed datasets is analyzed separately, and (iii) the *m* resulting estimates are combined using established procedures (31–33). The number of imputations was set to *m* = 20 for all methods. ***Runday-specific LOD-based methods (RC-R*/*ITS-R*/*MITS-R):*** The previously described methods *RC, ITS,* and *MITS* are applied within run days where at least 17 observations are available. In *RC-R*, the remaining missing values are set to the mean of all available expected values. For *ITS-R* and *MITS-R*, the remaining missing values are replaced using *ICE-norm* (see below). ***Imputation by chained equations (ICE-norm*/*-pmm*/*-adjR)*** was performed using the *R* package *mice*, version 2.25. It uses a repeated chain of equations through the incomplete variables, where in each imputation model, the respective incomplete variable is modeled as a function of the remaining variables (34–36). In *ICE-norm*, a Bayesian linear regression is used as the imputation model, whereas in *ICE-pmm* (predictive mean matching as imputation model), missing values are replaced by a random draw of measured values from other observations with the closest predicted values. In *ICE-adjR*, a model is specified with random intercept per run day, which aims to better utilize run day information. This model assumes that variable values (i.e., metabolite concentrations) have a run day-specific component, which varies randomly following a normal distribution. ***Multiple imputation by chained equations (MICE-norm/-pmm/-adjR)*** was performed using the *R* package *mice*, version 2.25: *MICE-norm, MICE-pmm,* and *MICE-adjR* consisted of *m* = 20 parallel imputation runs of *ICE-norm, ICE-pmm,* and *ICE-adjR,* respectively. Subsequently, the estimates are combined using Rubin’s rules as described above for *MITS*. ***MICE average version (MICE-avg-norm/-pmm)*:** *ICE-norm* or *ICE-pmm* is applied multiple (*m* = 20) times in parallel, followed by combining the *m* imputed datasets to one final dataset as the average of the imputed values. ***K-nearest neighbor imputation (KNN-var(K)/KNN-obs(K)/KNN-obs-sel(K)):*** In *KNN-var* and *KNN-obs*, missing values of each variable are replaced by the weighted average of pre-specified *K* nearest variables and observations, respectively. Distances to neighbors were defined as Euclidean distance and weights were chosen as *e*^*–d*^, where *d* defines the distances between two variables or observations. In *KNN-obs-sel, KNN-obs* is performed by selecting the strongest correlated variables with |*ρ*| ≥ 0.2, but it was constrained to a minimum of 5 and a maximum of 10 variables. The0020number of neighbors for *K* was set to 3, 5, 10, and 20.

More detailed descriptions of *RC*, *RC-R*, *ITS*, *MITS*, *ICE*, and *KNN*-based methods can be found in File S4. The two best performing methods, *KNN-obs-sel(K)* and *MICE* are available as R code in File S11.

### Statistical evaluation of missing data handling strategies in the simulation study

Pearson correlation, partial correlation, linear regression, and logistic regression analysis were performed, and the ability of imputation methods to reconstruct true associations and unbiased hypothesis test results was evaluated. For logistic regression, a dichotomized variable was simulated by discretizing one of the simulated continuous variables: all values above the median were set to 1 and all values below the median were set to 0. This dichotomized variable was used as response and the remaining continuous variable as predictor. For MI strategies, the resulting (correlation or regression coefficient) estimates and their variances were combined using Rubin’s rules. The obtained point estimates were then compared with the true underlying values by assessing the validity of hypothesis tests. To this end, type 1 error was calculated as the proportion of significant estimates (at α= 0.05) after imputation when there was no true effect. Power was calculated as the proportion of significant estimates (at α= 0.05) after imputation in the presence of a true effect. Detailed results can be found in File S5.

### Evaluation based on pathway modularity

This analysis was based on pathway annotations from Metabolon Inc. (see Supporting Information S9). Each imputation strategy was applied to the KORA F4 metabolomics data, resulting in different imputed datasets. All unknown metabolites were excluded since these compounds were not assigned to a pathway. For each imputed dataset, a Gaussian graphical model (GGM) was estimated to infer a network using the *R* package *GeneNet*, version 1.2.12. In previous studies, we have demonstrated that these models correctly reconstruct biochemical pathways from the data (22,25,37). In the case of MIs, a GGM was estimated for each imputed dataset, followed by combining partial correlations using Rubin’s rules after a Fisher Z-transformation. The network was constructed using partial correlations that are significantly different from zero after Bonferroni correction for *n* * (*n* – 1)/2, where *n* is the number of metabolites.

The pathway-based network modularity measure *Q* (22,24) was calculated for each network as

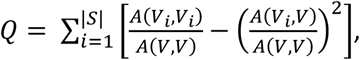

where |*S*| is the total number of pathways, *V* is the set of all metabolites, and *V*_*i*_ describes the subset of metabolites annotated with pathway *i*. *A*(*V*_*i*_, *V*_*j*_) is the number of edges between any two node sets *V*_*i*_ and *V*_*j*_. The variance of *Q* was estimated non-parametrically using bootstrapping of the original dataset (R package *boot*, version 1.3-15) with 1000 runs.

### Evaluation based on metabolite-SNP associations

Linear regression was performed using KORA F4 *CCA* and the results were compared with each other. For this analysis, we selected metabolite-SNP pairs for which (i) a genome-wide significant association could be identified in the meta-analysis of KORA F4 and TwinsUK cohorts in a previous GWAS (25) (summary statistics retrieved from http://www.gwas.eu); (ii) the proportion of each metabolite’s missing values in KORA F4 was between 10% and 70%; (iii) the metabolite was measured in the EPIC-Norfolk cohort, which we used to further benchmark the preservation of effect sizes; and (iv) a functional connection between the genetic locus of the SNP and the metabolite (e.g., metabolite is a known substrate of the transporter) was evident according to manual curation of the GWAS results (Table S8). For each imputed dataset, 18 metabolite-SNP pairs were tested for genetic association using age- and sex-corrected linear regression models under the assumption of an additive genetic model (metabolite ~ *β*_0_ + *β*_1_ × SNP + *β*_2_ × age + *β*_3_ × sex). To avoid spurious associations, metabolic data points greater than four SDs from the mean were removed prior to computing linear models. For MI approaches, the regression coefficients were pooled using Rubin’s rules as provided by the *R* package *mice*, version 2.25. For each metabolite-SNP pair, the variance of the regression coefficients and p-values were estimated using bootstrapping.

To explore which imputation approaches increased statistical power, p-values obtained for the effect sizes based on imputed data were compared with p-values obtained from *CCA* by calculating their ratio as 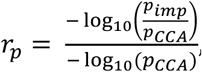, where *pimp* was the p-value obtained for imputed data and *p*_*CCA*_ was the p-value derived from *CCA*. A ratio less than or equal to zero indicated either no power gain or a power loss, whereas a ratio greater than zero indicated a drop in p-value, which suggested that statistical power increased when imputation was performed.

In addition to statistical power gain, the imputation approaches should be able to preserve effect sizes compared to *CCA*. Standardized effect sizes obtained from the imputed data (*β_imp_*) were compared with standardized effect sizes estimated for *CCA* (*β_CCA_*) based on the KORA F4 data (n = 1750) and the EPIC-Norfolk data (n = 10 634), assuming estimates from the EPIC-Norfolk data to be close to true effects. We calculated the ratio 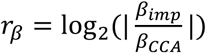, with a low ratio indicating a similar effect size between the imputed data and *CCA*. A highly negative or positive *r*_*β*_ indicates an underestimation or overestimation of the effect sizes in imputed data, respectively. A well performing imputation method is assumed to obtain high *r*_*p*_ and low absolute *r*_*β*_.

**Supporting information captions**

File S1. LOD tendency.

File S2. Runday-dependent densities in relation with missingness.

File S3. Simulation framework.

File S4. Imputation methods.

File S5. Simulation evaluation results.

File S6. Metabolite-SNP associations–beeswarm plots.

File S7. Metabolite-SNP associations compared with EPIC-Norfolk.

Table S8. Metabolite-SNP associations–linear regression results.

Table S9. KORA F4 annotations.

File S10. KORA F4 experimental setup.

File S11. KNN-obs-sel and MICE imputation code.

